# Bee phylosymbiosis: the relative importance of host phylogeny, behavioral traits, and environment

**DOI:** 10.64898/2026.10.19.760025

**Authors:** Linyang Sun, Johan Ekroos, Anne Duplouy

**Affiliations:** Department of Organismal and Evolutionary Biology, Faculty of Biological and Environmental Sciences, University of Helsinki, Helsinki, Finland; Faculty of Agriculture and Forestry, University of Helsinki, Helsinki, Finland

**Keywords:** Host-microbe association, Metabarcoding, Wolbachia, Pollinators, Biogeography

## Abstract

Phylosymbiotic studies investigate how microbiomes recapitulate their host phylogeny and can thus reveal the evolution of host-microbe relationships. In bees, the extent of phylosymbiosis and its relationship with host ecological traits remain unclear. We integrated public bee 16S rRNA metabarcoding data to investigate how host phylogeny and ecological traits affected the bacterial community associated with 178 bee species. Our results highlight only moderate phylosymbiosis patterns in bees, with the corbiculates associating with diverse lineage-specific microbial communities, while the non-corbiculates were predominantly associated with *Wolbachia*. Across all bee taxa, *Wolbachia* was also consistently negatively associated with *Lactobacillus*. Furthermore, our analysis revealed that the host ecological traits had only marginal effects on microbial composition. Although eusocial bees harbored distinct microbiomes, the effect of social behavior was confounded by host phylogeny. Climate showed no general effect across bee taxa, but within some genera, microbiomes were more homogeneous among bees from arid and semi-arid regions than among those from other climatic regions. Together, our findings reveal that broad-scale patterns in bee microbiomes emerge primarily as lineage-specific host–microbe associations, with ecological traits acting only weakly and often at a taxon-specific level.

## Introduction

Patterns in phylosymbiosis reflect host-microbe evolutionary associations in which microbial community composition mirrors host phylogeny [1], potentially arising through ecological filtering by host traits [2]. Such evolutionary patterns have been well described in the corbiculate bees [3]. However, whether phylosymbiosis is broadly evident across the entire bee phylogeny and their diverse ecological traits remains unclear as most studies have only examined few bees from limited geographic regions. The recent growing interest in host-microbe interactions has fueled the accumulation of microbial data from numerous bee species, making analyses possible across the entire clade of these important pollinators.

Bees are known to acquire microbes from the flowers they visit, the environments they are exposed to, and their nestmates during social interactions [4]. These acquisition routes are strongly linked to the ecological traits of bees. As an example, shorter-tongued bee species that frequently visit shallow flowers are associated with a high microbial diversity [5], and some microbes originating from flowers visited contribute to an increased nutritional status of the larvae of *Osmia ribifloris* [6].

Solitary bees acquire their microbes predominantly from environmental sources [7], in contrast to eusocial bees, which in turn transmit a conserved core gut microbiota among nestmates [3, 8]. Bee-associated microbial species can also include maternally inherited symbionts, which can vary in abundance in bees under different conditions, and between bee castes. For example, while the abundance of the symbiotic bacterium *Sodalis* differs among social castes in halictid bees [9], the prevalence and abundance of the symbiotic bacterium *Wolbachia* vary between populations of solitary bees [10] exposed to different local thermal conditions [11]. In addition to bees, experimental studies in *Drosophila melanogaster* have shown that infection with *Wolbachia* affects the abundance of *Acetobacter pasteurianus* in the flies [12]. Thus, the presence of symbiotic bacteria can have important effects on the overall composition of the host-associated microbiota. However, such patterns remain to be investigated in bees.

Several ecological traits are however constrained by phylogeny, with closely related bee species showing more similar traits [13]. For example, advanced eusociality is restricted to the corbiculate bee clade [14]. Similarly, many recently diverging species or genera might still be sympatric and might utilize rather similar food resources and nesting substrates, and hence they might also acquire similar environmental microbes [15, 16]. As a result, the effect of host phylogeny on the composition of the bee microbiome is likely confounded with the effects of various bee ecological traits, such as social behavior, diet, nesting substrate, and climate. Although clade- or region-specific studies have already provided insight into the independent contribution of phylogeny versus diverse bee-traits, a global analysis spanning a broader range of bee diversity would provide greater power to disentangle this.

To address this gap, we assembled a global dataset of published bee 16S rRNA gene sequences, comprising 189 bee taxonomic units. This broad taxonomic and geographic coverage extends the study of phylosymbiosis beyond the few already intensively studied bee lineages [3] and enables us to examine whether host– microbiome associations are consistent across diverse bee ecological and behavioral contexts. By examining phylosymbiosis across this broader diversity of bees, we better identify the ecological and evolutionary factors that shape their microbial associations.

## 2 Material and Methods

### 2.1 Data compilation and metadata curation

We used the Meta2Data, described by Li et al. [17], to assemble a wild bee metabarcoding dataset. The datasets were retrieved from four major public repositories (ENA, CNCB, NCBI, and DDBJ), with a combination of keywords related to typical methodological approaches (‘amplicon’, ‘16S’, ‘metabarcoding’, ‘bacteria’, ‘microbiome’, ‘microbiota’, and ‘microorganisms’) and to the host identity (“bee”, and each of the genus names from the bee phylogeny by Henríquez-Piskulich et al. 2024 [18]). All 16S rRNA amplicon sequencing datasets published before March 2026 were considered.

We recovered ecological metadata for each host species, including information on social behavior, nesting, and the degree of foraging specialization (lectism), from either the original study, or from diverse trait databases [19–22] (Supplementary doc1). These traits are highly associated with microbiomes. To estimate the effect of climate, we collected the geographic information of each sample. We only had access to the exact geographic coordinates for about 40% of all samples. For the rest, we retrieved locality information (i.e. at best region, or county) from the original publications and assigned coordinates based on the reported area, which is sufficient given that we focus on broad-scale climatic effects. We extracted 19 bioclimatic factors described by CHELSA V2.1 [23] and the Köppen-Geiger system [24] (1981-2010, ∼1 km resolution) from the geographic coordination. To reduce dimensionality and collinearity among the bioclimatic variables, we then summarized the CHELSA bioclimatic variables by principal component analysis, retaining only the first two axes. The first axis (50% of the variance) described a gradient from warm, wet, and weakly seasonal climates to cold, dry, and strongly seasonal climates. The second axis (18%) captured summer warmth and precipitation.

To conduct the most balanced comparisons, we applied sample-level filters to exclude sample types that are underrepresented in datasets. Similarly, we included only wild bees or samples from hives managed under natural conditions, but excluded any experimental treatment groups, such as antibiotic treatment, dietary manipulation, or pathogen infection. Because the developmental stages and castes can affect the microbiome composition in bees [25, 26], we also excluded data collected from larvae, pupae, and queens. We only included data from samples based on the entire gut or body, and discarded data from non-gut tissues or incomplete gut samples.

### 2.2 Sequence data preprocessing

The dataset included samples that were amplified by different primer sets, samples that covered different hypervariable regions, and samples sequenced on different platforms (e.g., Illumina, PacBio, Ion Torrent, Roche 454). We processed all raw sequencing data with the VSEARCH [27], as implemented in the Meta2Data pipeline [17]. We thereafter aligned the representative sequences generated by VSEARCH, and inserted them into a reference full-length 16S phylogenetic tree using the SEPP tool [28] to produce a bacterial phylogeny tree. Finally, we assigned taxonomy using a full-length 16S classifier trained on the SILVA 138.1 reference database [29].

### 2.3 Data analyses

To measure the community structure, we calculated the Hill diversity numbers, corrected for sequencing depth, using the R package iNEXT [30]. We obtained three corrected Hill numbers: q0 (richness), q1 (the exponential of Shannon entropy), and q2 (the inverse Simpson concentration). We removed all samples with an estimated q0< 3, as we considered that data to be the result of potential methodological flaws, and likely not representative of the true bacterial diversity within the samples. After all these cleaning steps, our dataset included a total of 5,218 samples, and 2,936 bacterial genera. We measured Beta diversity using Bray-Curtis dissimilarity, weighted and unweighted UniFrac, and Aitchison distance metrics. We used the Uniform Manifold Approximation and Project (UMAP) [31] to cluster samples based on Aitchison distance with 15 neighbours and min_dist = 0.1.

To assess batch effects, we built marginal PERMANOVA models (999 test) between metadata variables (BioProject, amplified 16S region, and sequencing platform) and Aitchison distances for the communities. To test the confounding effects between each variable, we also quantified the strength of each association using both standard and bias-corrected Cramér’s V, considering every pair of variables (BioProject, amplified region, platform, primer, tissue, sex, month, host species, host genus and host family).

#### 2.3.2 Variance partitioning

To partition alpha diversity according to ecological factors, we fitted separate Bayesian phylogenetic mixed models for each log-transformed Hill number using MCMCglmm [32]. The fixed effects included in the models were host sociality, lectism, nesting strategy, and PC axes (PC1 and PC2) describing the climate. To distinguish between-species and within-species associations between climate and alpha diversity, climate PCs were decomposed into two components: the host-species mean, and the deviation of each sample from its corresponding host-species mean (Mundlak decomposition). The species mean therefore represents the between-species component, whereas the sample-level deviation represents the within-species component. To account for non-independence among species and other sources of structured variations (i.e., batch effect), we included host phylogeny, 1° geographic grid-cell, and the Dataset ID (BioProject) as random effects. Models were fitted to 4,427 samples from 141 host species, and for 23,000 iterations, with a burn-in of 3,000 and a thinning interval of 20 iterations. We quantified the contribution of each fixed-effect group using variance partitioning of the linear predictor.

To assess the robustness of the variance estimates to each host taxa or each dataset, we repeated the analysis using the leave-one-out approach. Because of the computational cost of these 642 refitted models (i.e., leave-one-host-species-out and leave-one-BioProject-out), the leave-one-out analyses were run for 13,000 iterations, with the first 3,000 iterations discarded as burn-in and every 10th iteration retained, resulting in 1,000 posterior samples per refitted model.

We also measured the contribution of the different ecological factors to the beta diversity of the bee microbiota using sequential PERMANOVA (adonis2, by = “terms”, with 999 permutations). The community data was aggregated by BioProject x host species x 1° geographic grid cells to reduce the overrepresentation of densely sampled groups. For this analysis, we removed the bacterial genera with mean relative abundance below 0.1% across aggregated units. Host phylogeny was represented by principal-coordinates derived from the host patristic-distance matrix.

We only kept the axes that explained at least 80% of the variance associated with positive eigenvalues. The fixed terms were ordered as follows: host phylogeny, BioProject, sociality, lectism, nesting and climate. The reported R^2^ values represent the incremental variance explained by each term conditional of each preceding terms in the model, rather than independent marginal effects.

Host phylogenetic relatedness was represented by principal-coordinate axes derived from the host patristic-distance matrix, retaining sufficient axes to explain at least 80% of the variance associated with positive eigenvalues. We fitted sequential PERMANOVA models with the terms entered in the following order: host phylogeny, BioProject, sociality, lectism, nesting strategy, and climate. Each model was evaluated using 999 permutations. The reported R² values therefore represent the incremental variance explained by each term conditional on the preceding terms in the model, rather than independent marginal effects.

#### 2.3.4 Phylosymbiosis analysis

To assess patterns of phylosymbiosis in bees, we applied complementary abundance and occurrence-based approaches. The abundance-based analysis was performed using ABDOMEN [33], a Bayesian multivariate Brownian-motion model that evaluates the correspondence between microbial community composition and host phylogeny. We agglomerated the microbial abundances at the bacterial order level and converted them to relative abundances. Bacterial orders with a mean relative abundance of at least 0.1 % in at least one host species were retained, while all remaining and unclassified orders were pooled in a ‘Others’ category. Samples within each host species were then averaged to obtain a representative microbial community per host species. The ABDOMEN model fitted with a detection threshold of 1e-5, an empirical root-state prior, 20 chains, 1,600 total iterations per chain, and 1,000 warm-up iterations.

For the occurrence-based analysis, performed on the host-species phylogeny, we followed a framework established for mammalian gut microbiota [34] and corbiculate bees [3]. Only the host species represented by at least five samples were included, and any bacterial genus was considered present in a sample when its relative abundance exceeded 0.1%. For each host species, the prevalence of each bacterial genus was calculated as the proportion of samples in which the genus was detected. To reduce potential false-positive associations caused by low-prevalence taxa, the bacterial genera were used in the ancestral-state analysis only when their prevalence reached at least 50% in at least two host species. Host species were only considered carrying a particular bacterial genus, when the prevalence of this genus was at least 30%. We selected the 30% threshold, after evaluating prevalence thresholds ranging from 10% to 70%.

For each bacterial genus, we fit alternative models and selected the best-fitting model using AICc. Ancestral-state probabilities were reconstructed using phytools [35], and the reconstructions were used to estimate lineage-specific gains and losses of each bacterial genus along the host phylogeny. To test whether the distribution of the bacterial genera across the host species exhibited significant host-phylogenetic structure, we performed 499 tip-state permutations. The resulting p-values were then corrected for multiple testing across genera using Benjamini–Hochberg corrections. All retained genera were clustered according to their host-species presence patterns, to identify genera showing similar phylogenetic distributions.

#### 2.3.5 Microbial community associations

To estimate the relationship among bacterial genera across host groups, we first inferred microbial association networks using SparCC [36]. These networks were constructed separately for all bees, corbiculate bees only, and non-corbiculate bees only, as these groups showed distinct patterns in previous analyses. To reduce unstable associations caused by rare bacterial taxa, and to improve the robustness of network inference, we included only bacterial genera detected in at least 20% of the host samples. The bacterial symbiont *Wolbachia* was retained regardless of its prevalence because of its biological importance, this despite its relatively low abundance in most corbiculate bee samples. The SparCC correlations were estimated using 999 bootstrap replicates, and P-values were corrected for multiple testing using the Benjamini-Hochberg correction test across all genus pairs within each network.

To further quantify the association between *Wolbachia* and corbiculate associated genera, we fitted CLR-based linear regression models separately for corbiculate and non-corbiculate hosts. To account for the non-independence of samples from the same host species, uncertainty was assessed using a host-species block bootstrap. Samples were included in the analysis only when both *Wolbachia* and the second focal bacterial genera had at least five reads. CLR-transformed abundance values were calculated using a pseudo count of 0.5. Host species were resampled for 20,000 bootstrap replicates. Finally, we summarized *Wolbachia* occurrence across host genera using all samples to characterize the distribution of this symbiont among host taxa.

## 3 Results

### 3.1 The global bee microbiome dataset

After curation, our final dataset included 5,218 samples from 83 datasets, representing 189 bee taxonomic units. Of these taxonomic units, 178 (94%) were identified to species level and accounted for 5,035 samples (96.5%), while the remaining 11 genus level units (5.8%) included 183 samples (3.5%). Apidae accounted for 82.8% of the samples in the dataset, with the *Apis* and *Bombus* genera being the most represented (35.7% and 23.3% of all samples, respectively). *Apis mellifera* alone accounted for 29.5% of the samples (Table S3). Sampling was geographically uneven and concentrated in the Northern Hemisphere. Together, North America and Europe contributed 46.6% of all samples (Figure 1A; Table S1).

**Figure 1:**
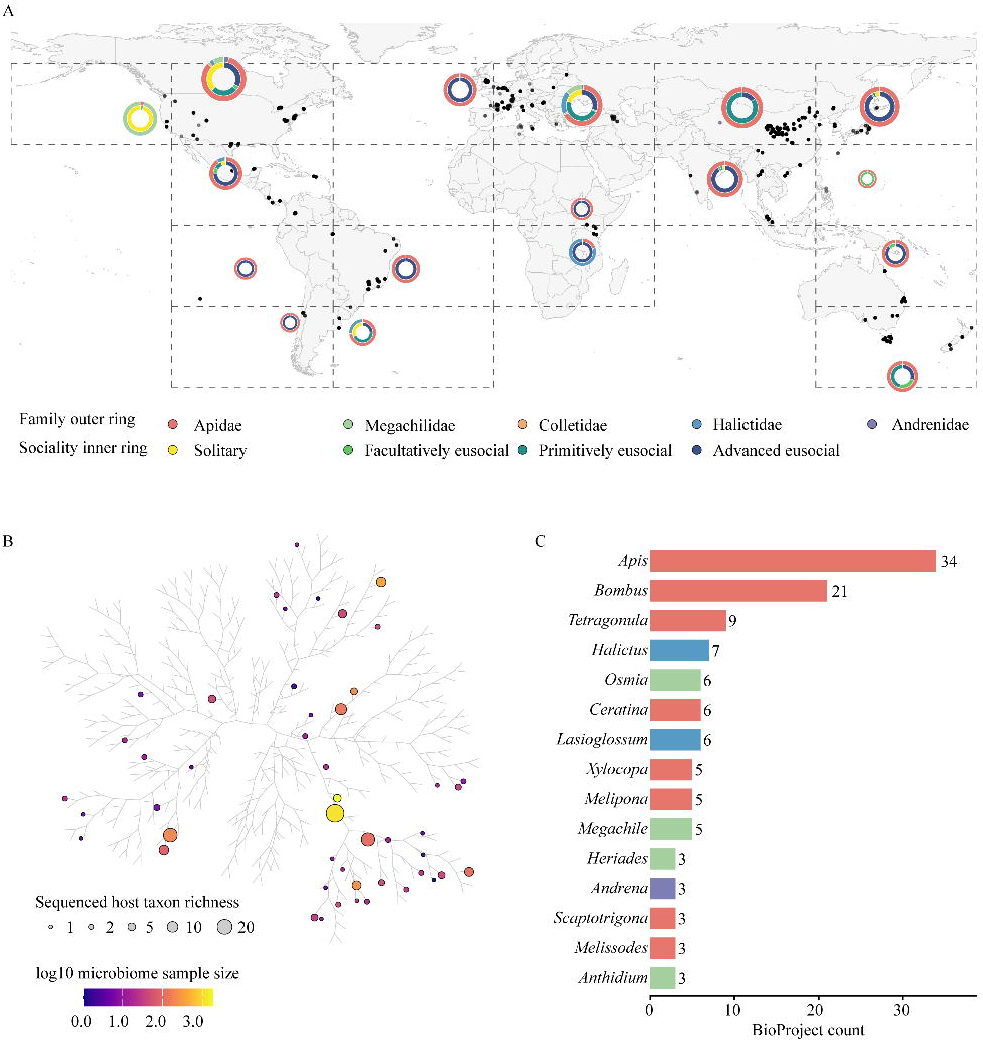
Global and phylogenetic coverage of the bee microbiome dataset is extensive but taxonomically uneven. The final dataset comprised 5,218 bacterial-community samples from 189 operational host taxonomic units (178 species-level units), five families, and 83 BioProjects after applying a minimum sequencing-depth threshold of 2,500 reads. (A) Global sampling distribution. Black points show each of geographic coordinates. Dashed rectangles delimit the 17 occupied 60° longitude × 30° latitude grid cells. For each occupied cell, the outer donut ring shows the proportions of samples from Apidae, Halictidae, Megachilidae, Andrenidae, and Colletidae, whereas the inner ring shows host sociality. Donut diameter increases with the number of samples in the cell. (B) Genus-level bee phylogeny showing microbiome sampling coverage. Grey branches represent all 427 genera in the reference tree. The circles mark the sampled genera represented in the tree. Circle area scales with the number of sequenced host taxonomic units, and fill color represents the log10-transformed number of microbiome samples. (C) The 15 most intensively studied bee genera ranked by the number of BioProjects. Bar colors show host family, and numbers at bar ends give BioProject counts.

We also report technical biases in the dataset. Our datasets include samples from 11 different hypervariable regions, of which V4 was the most amplified (42.0%). Most samples were sequenced on Illumina platforms (86.1%; Tables S1 and S2). After quality controls, sequencing depth ranged from 2,504 to 1,315,721 reads, with a median of 27,866 reads, and a mean of 46,436 reads (Figure S1; Table S4). Among 2,936 microbial genera detected, 61.0% of the reads were assigned to bacterial genera commonly found in corbiculate bees, including *Lactobacillus* (11.97%), *Gilliamella* (11.68%), *Snodgrassella* (10.93%), and *Bombilactobacillus* (5.09%). The ten most abundant bacterial families, including Lactobacillaceae (23.14%), Orbaceae (16.39%), Neisseriaceae (13.31%), and Bifidobacteriaceae (6.42%) (Table S5), accounted for 81.5% of the mean within-sample relative abundance. The bacterial genera *Apilactobacillus* and *Acinetobacter* were the only two to occur across all five collected bee families accounted for in our study.

### 3.2 Variance partitioning between phylogeny, behavior traits, and climate

Bacterial community composition varied markedly among BioProjects. In a marginal PERMANOVA, BioProject explained 41.7% of the variation in the Aitchison distance (R² = 0.417, P = 0.001; 999 permutations; Table S6). BioProject was however strongly confounded with host species: 144 of 177 host species (81.4%) occurred in only one BioProject in the dataset used to assess batch effects (n = 4,870 samples). Host species were also strongly associated with the 16S region amplified (Cramer’s V = 0.793; bias-corrected V = 0.770; n = 5,051; Figure S2; Table S6).

#### 3.2.1 Variance partitioning for alpha diversity

The genus-level Hill diversity varied substantially (Figure 2A). Of the three Hill orders, only q0 showed a weak association with host-genus phylogenetic distance (Mantel test, r = 0.249, BH-adjusted P = 0.029; 5,000 samples, 34 genera; Table S7). This indicates that differences in bacterial richness (at the genus-level) tended to increase with phylogenetic distance between host species.

**Figure 2:**
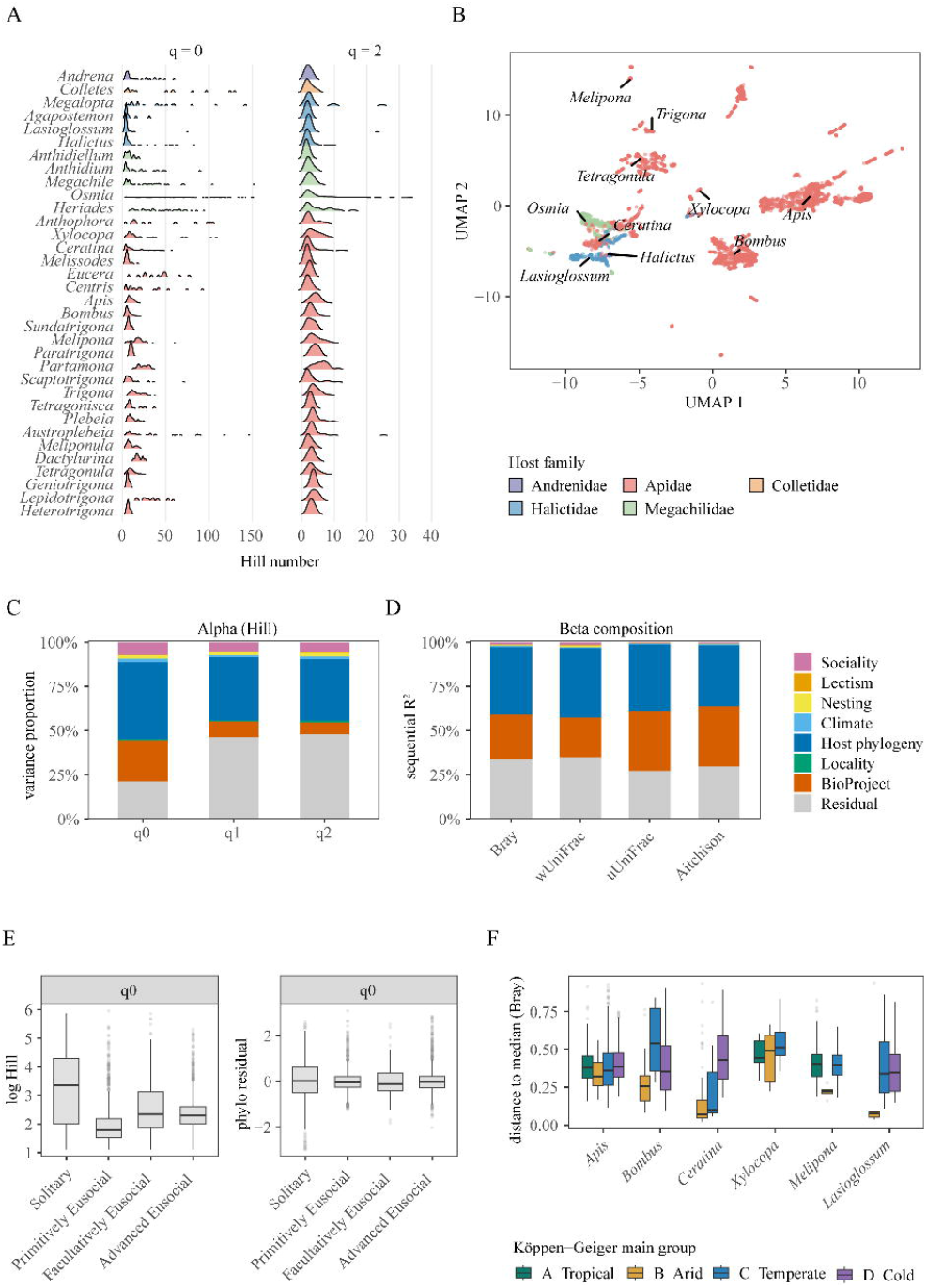
Host phylogeny accounts for most variation observed in bee-associated bacterial diversity and composition. (A) Alpha diversity density plot across 34 bee genera represented by at least 10 samples. Genera are ordered according to host phylogeny, and ridge colors indicate host family. Hill diversity is shown for q = 0 and 2. (B) Uniform Manifold Approximation and Projection (UMAP) embedding of all 5,218 samples. Each sample is colored by host family. The italic labels mark the median UMAP coordinates of the ten host genera with the largest sample sizes. (C) The mean variance proportions for log-transformed Hill diversity at q = 0, 1, and 2 from phylogenetic mixed models. Fixed-effect contributions comprise sociality, lectism, nesting, and climate. (D) Sequential variance partitioning of bacterial community composition for Bray–Curtis, weighted UniFrac (wUniFrac), unweighted UniFrac (uUniFrac), with terms entered in the order of host phylogeny, BioProject, sociality, lectism, nesting, and climate. (E) Distribution of log q = 0 Hill diversity among host-sociality classes before (left) and after (right) removal of the host-phylogenetic component (5,010 samples from 169 species). (F) Within-genus distance to the multivariate median of each Köppen–Geiger main climate group, calculated from Bray–Curtis dissimilarities for six well-sampled host genera existed in three or more climate groups (tropical; arid; temperate; or cold).

The variance partitioning was based on 4,427 samples from 141 host species across 73 BioProjects. After accounting for other ecological factors, host phylogeny accounted for the largest proportion of variance, ranging from 43.4 for q0 and 35.0% for q2. In contrast, BioProject accounted for 23.3%, 8.8%, and 6.9% of the variance for q0, q1, and q2, respectively. Among the ecological trait predictors, sociality explained the largest proportion of variance (5.2–7.1% of each q-value), whereas each of the remaining ecological factors only accounted for less than 2% independently (Figure 2C; Table S8).

The q0 distribution changed substantially after accounting for host phylogeny. Across the sociality categories, the median raw log-transformed q0-values ranged from 1.79 to 3.35, whereas median phylogenetic residuals ranged from −0.119 to 0.016 (Figure 2E). Hence, a substantial proportion of the variation in observed richness among sociality categories was associated with the structure of bee phylogeny.

Finally, the variance estimates for all three Hill orders were robust to the exclusion of any single host species or BioProject. The leave-one-out analyses revealed no evidence that the estimated contributions of ecological factors were driven by any single host species or study (Table S9).

#### 3.2.2 Variance partitioning for beta diversity

A CLR-based UMAP of all 5,218 samples showed pronounced host taxonomic structure in bacterial community composition (Figure 2B). Recolouring the same coordination by climate, sociality, and nesting strategy revealed additional patterns that were largely overlapped with those associated with host taxonomy (Figure S3).

The sequential PERMANOVA was performed on 396 trait units, each defined by a unique combination of BioProject, host species, and rounded 1° geographic grid cells (Figure 2D; Table S8). Across the four-distance metrics, host phylogeny accounted for 34.6 to 39.3% of the variation, while BioProject accounted for 22.5 to 34%. In contrast, the ecological factors each explain relatively small fractions of variation (i.e., sociality: 0.4–1.5%, climate: 0.5–0.7%, nesting: 0.2–0.7%, and lectism: 0.2–0.5%).

Because climate effects might be obscured by differences among host species, we further compared the bacterial composition of six host genera well sampled across climatic zones. Community composition was more homogeneous in arid climate regions, as indicated by smaller Bray–Curtis distances to the corresponding Köppen climate-group median, whereas temperate regions showed greater within-group heterogeneity. (Figure 2F).

### 3.3 Pattern of phylosymbiosis in bees

The abundance-based ABDOMEN analysis suggested a moderate phylogenetic signal in bacterial community composition across the bee phylogeny (Pagel’s λ = 0.368, 95% credible interval [0.317, 0.421]; Figure 3; Table S10).

**Figure 3.**
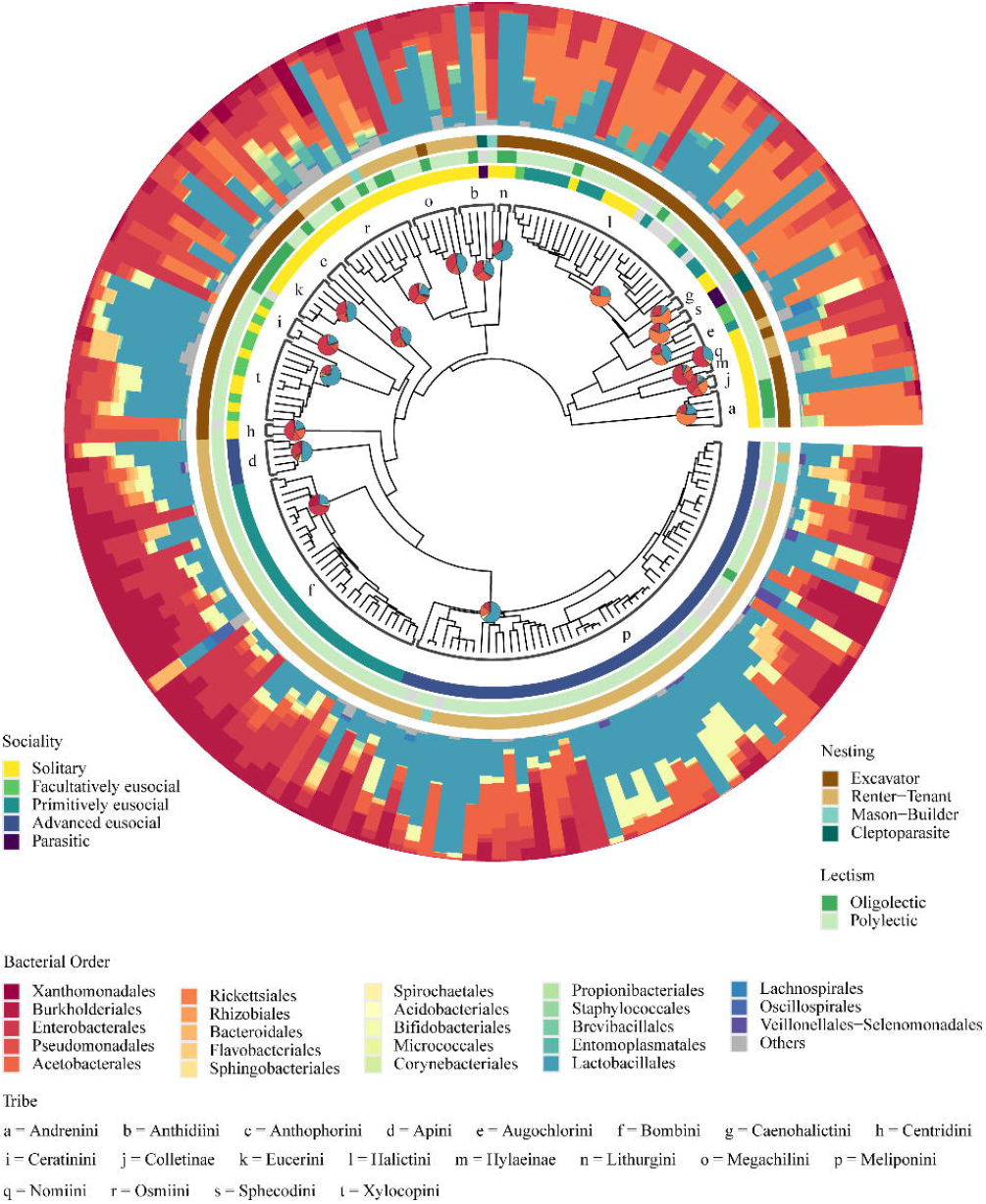
Phylogenetic reconstruction of order-level bee microbiome composition across host lineages. The circular composite summarizes bacterial order-level profiles for 5,035 samples representing 178 host species. The central fan tree is the host-species phylogeny. The lowercase letters identify 20 host taxonomic groups. The pies at the most recent common ancestor show ancestral bacterial-order compositions estimated with ABDOMEN. From the tree outward, the three narrow rings denote host sociality, lectism, and nesting strategy, respectively, using the colors defined in the keys, and grey indicates missing trait information. The outer stacked bars show the mean relative abundance of bacterial orders for each host species. Orders reaching at least 10% mean relative abundance in one or more host species are shown separately, whereas the remaining orders are pooled as “Others.” Across the 178 host species, the estimated global phylogenetic signal was Pagel’s λ = 0.37 (95% credible interval, 0.32–0.42).

The occurrence-based model showed that phylogenetic structuring was restricted to a small subset of bacterial taxa. Of 138 bacterial genera tested across 111 host species, 51 showed a significant phylogenetic signal after false-discovery-rate correction (FDR-adjusted q < 0.05). After excluding 21 unclassified genera and one rare genus, 29 genera were used for clustered in four modules based on their presence/absence patterns across the host phylogeny (Figure 4; Table S10). The largest module (m2) comprised 18 bacterial genera, including *Snodgrassella*, *Gilliamella*, *Bombella, Commensalibacter*, *Apibacter, Bifidobacterium, Lactobacillus,* and *Bombilactobacillus*. These bacterial genera had higher mean prevalence across in corbiculate than in non-corbiculate host species (36.3 vs. 15.1%, respectively) (Figure 4; Table S10). In contrast, bees from the Megachilidae family consistently harbored a distinct group of bacteria, belonging to the module m3. The bacterial symbiont *Wolbachia* occurred in only 5.4% of the corbiculate species (N=3/56), but in 52.7% of the non-corbiculate species (N=29/55) (Figure 4; Table S10).

**Figure 4.**
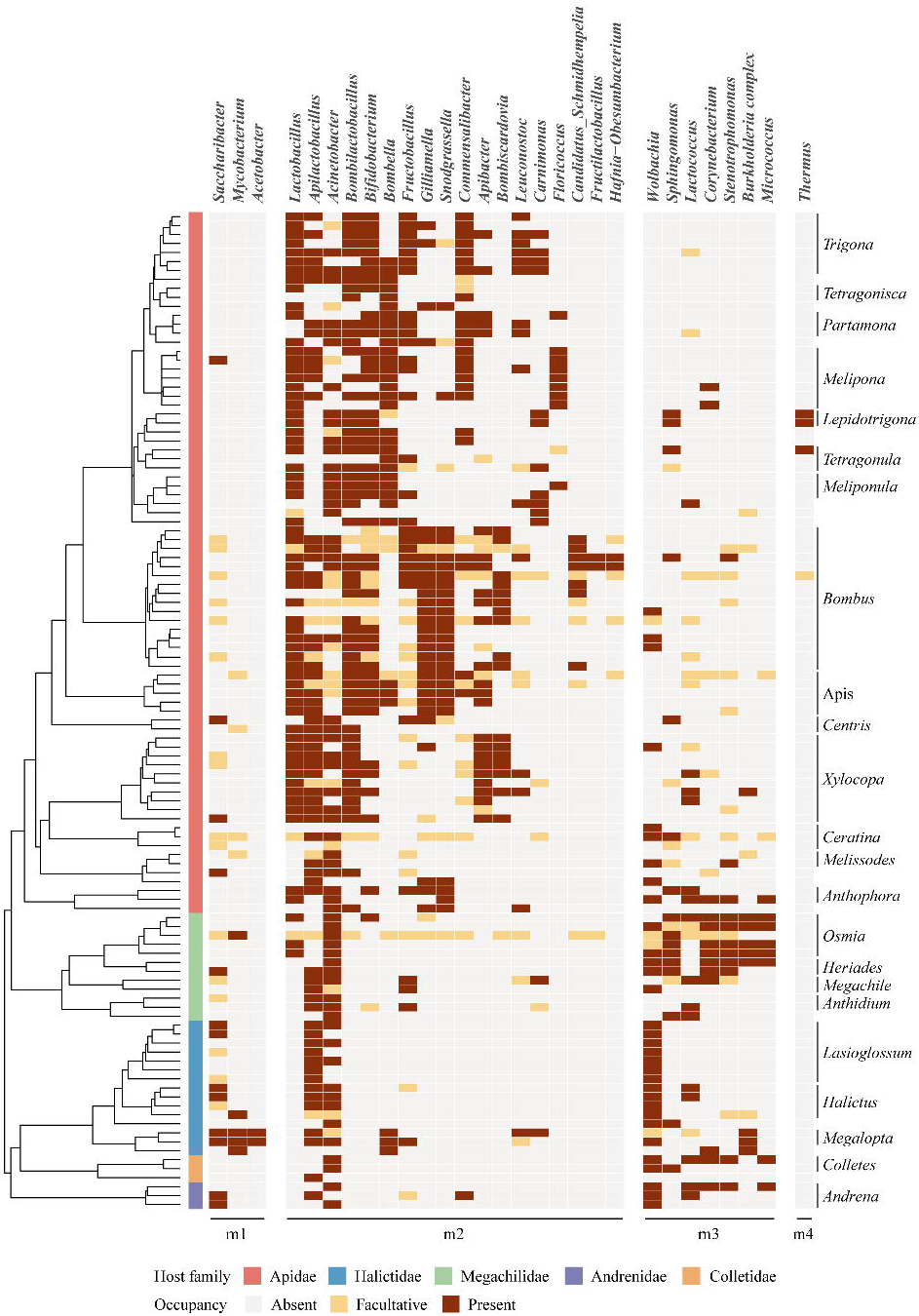
Phylosymbiosis pattern across bees based on occurrence-based model. The figure shows the phylosymbiosis pattern in bee included in 111 bee species represented by at least five samples.The color band shows host family, and the heatmap shows bacterial genus occupancy. The matrix includes 29 bacterial genera with significant host-phylogenetic structure based on a 499-permutation tip-shuffle test (Benjamini–Hochberg-adjusted q < 0.05), grouped into four co-occurrence modules (m1–m4). Host genera represented by at least two species are labelled on the right.

### 3.4 Microbial associations with *Wolbachia*

The pooled SparCC network included 5,218 samples and 29 bacterial genera. We found ten significant associations between *Wolbachia* and other bacteria (|r| ≥ 0.20, BH-adjusted q < 0.05; Figure 5A; Table S12). Negative associations were observed with *Lactobacillus*, *Bombilactobacillus*, *Bifidobacterium*, and *Commensalibacter* (r = −0.32 to −0.24), whereas positive associations were detected with *Pseudomonas*, *Acinetobacter*, *Methylobacterium–Methylorubrum*, *Apilactobacillus* (r = 0.27–0.33). The associations between *Wolbachia* and other bacterial genera strongly differed between host-clades. We observed one positive association between *Wolbachia* and *Bombiscardovia* (r = 0.33) in the corbiculate bee network, while the non-corbiculate network showed one negative associations between *Wolbachia* and *Methylobacterium–Methylorubrum* (r = −0.21), and positive associations with *Streptococcus*, *Stenotrophomonas*, *Saccharibacter*, *Apilactobacillus*, and *Sodalis* (r = 0.20–0.57; Figure 5A; Table S12).

**Figure 5:**
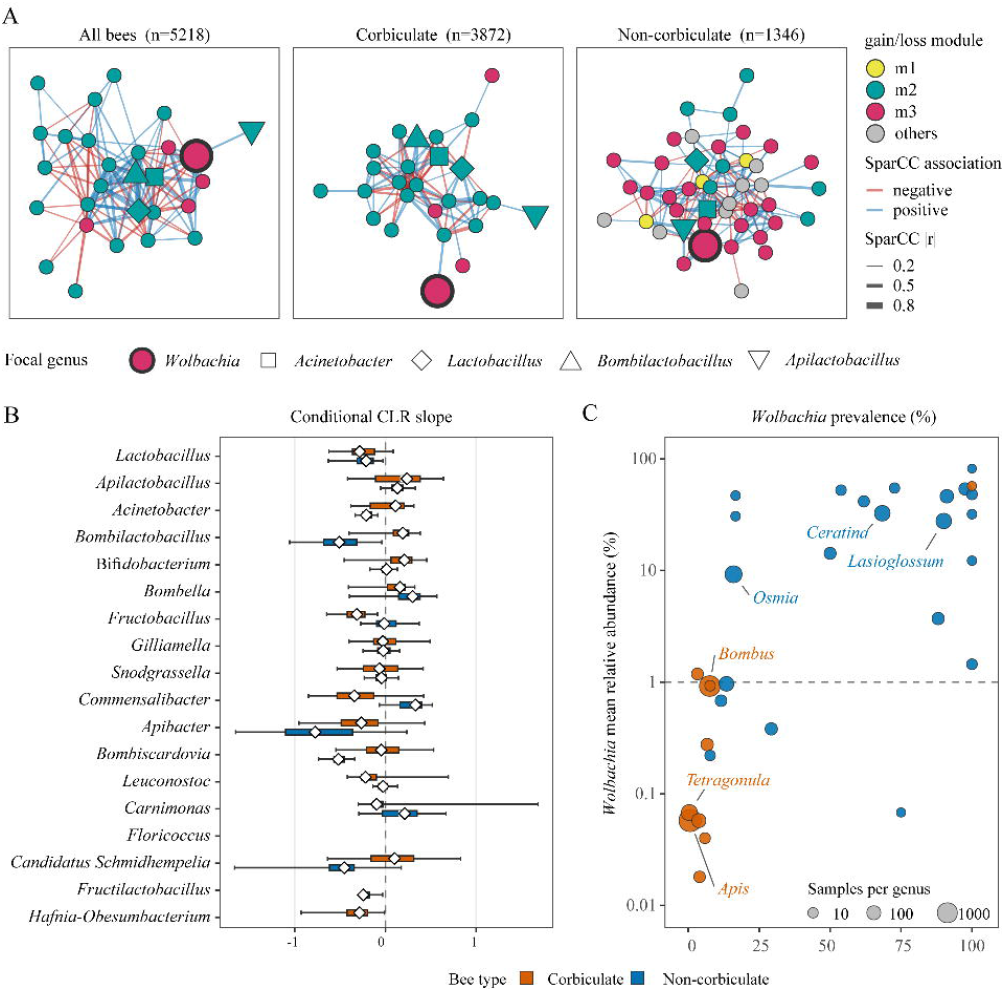
***Wolbachia* prevalence and bacterial-community associations show clade-specific patterns in corbiculate and non-corbiculate bees.** (A) Sparse Correlations for Compositional data (SparCC) genus-level association networks for all bees (n = 5,218), corbiculate bees (n = 3,872), and non-corbiculate bees (n = 1,346). Only the genera detected in at least 20% of the samples were analysed (up to the 80 most prevalent genera). Blue and red links denote positive and negative SparCC correlations, respectively. Displayed links satisfy |r| ≥ 0.20 and Benjamini– Hochberg-adjusted q < 0.05 based on 999 bootstrap replicates. Node fill identifies the gain/loss module. Nodes are shaped according to microbial genera: Wolbachia (red circle), Acinetobacter (square), Lactobacillus (diamond), Bombilactobacillus (upward triangle), and Apilactobacillus (downward triangle) and non-focal genera are normal size circles. Node size distinguishes Wolbachia, the other focal genera, and non-focal genera. (B) Bivariate associations between *Wolbachia* and the 18 m2 genera. The x axis gives the slope of partner-genus CLR abundance on *Wolbachia* CLR abundance. Slopes were estimated separately within each host clade. The vertical dashed line denotes a slope of zero. (C) Host-genus *Wolbachia* prevalence versus mean relative abundance among positive samples, using a positivity threshold of at least five reads across all 5,218 samples. Each point represents a host genus. Orange denotes corbiculate hosts, and blue denotes non-corbiculate hosts. The point size denotes the total number of samples in that host genus. The horizontal dashed line marks 1% mean relative abundance.

*Wolbachia* was detected in 16.0% of all the bee samples, with only 3.6% of the 3,872 corbiculate samples being infected, and 51.5% of the 1,346 non-corbiculate samples carrying the bacterium. By examining the associations between *Wolbachia* and the 18 other bacterial genera from module m2 using CLR-based regression with host-species block bootstrap, we showed that out of the 36 clade-by-genus combinations, 32 (88.9%) met our criteria for sample-size, host-species-blocking, slope-estimation, and bootstrap estimation. Out of these 32, we found that 21 (65.6%) complete-data slopes were negative and 11 (34.4%) were positive. Seven negative associations had 95% bootstrap intervals below zero. In non-corbiculate hosts, these associations involved *Lactobacillus* (β = −0.212, 95% interval [−0.628, −0.027]), *Acinetobacter* (−0.212 [−0.333, −0.084]), *Bombilactobacillus* (−0.509 [−1.055, −0.040]), *Bombiscardovia* (−0.518 [−0.737, −0.337]), and *Fructilactobacillus* (−0.244 [−0.289, −0.031]). In corbiculate hosts, the negative association involved *Fructobacillus* (−0.315 [−0.648, −0.087]) and *Hafnia–Obesumbacterium* (−0.285 [−0.928, −0.011]). The remaining 25 eligible combinations showed varying degrees of directional support, although their 95% bootstrap intervals overlapped zero. Notably, in the corbiculate hosts, the *Lactobacillus* bacteria showed a negative point estimate (β = −0.284, 95% interval [−0.623, 0.085]), with 92.2% of bootstrap slopes below zero, suggesting a predominantly negative association but higher uncertainty (Figure 5B; Table S12).

*Wolbachia* was detected in 32 of the 50 bee genera (using the five-read detection criterion), whereas 18 host genera, representing 332 samples, were *Wolbachia*-free (Figure 5C; Table S13). Among the 32 positive host genera infected with *Wolbachia,* the prevalence of the bacterium was positively correlated with its mean relative abundance among positive samples (Spearman ρ = 0.742). Together, these results indicated a pronounced host-clade dependence in *Wolbachia* prevalence and its associations with other bacterial genera. These network and CLR-based analyses identified conditional co-occurrence patterns and associations, but do not demonstrate direct microbial interactions or causal relationships.

## 4. Discussion

Our study represents a highly comprehensive re-analysis of wild bee microbiome data, drawing upon 189 bee taxonomic units from 83 independent studies. We showed that after accounting for host phylogeny, the different bee behaviors (sociality, nesting, lectism) explained only little of the variation in alpha and beta diversity in their associated bacterial communities. Climate had also no consistent overall effect across all bees, however, within some broadly distributed genera, bees from arid and semi-arid environments tended to have more similar microbiomes than bees from other climatic zones. Phylosymbiosis is associated with an Apidae-associated bacterial community and the presence of the endosymbiotic bacterium *Wolbachia*, common in many other insects [37]. These two communities showed an almost inverse distribution: where the Apidae-associated community was observed, *Wolbachia* tended to be scarce, and vice versa. The Apidae-associated community was common in the corbiculate clade, as well as in *Xylocopa* and *Centris* bees, with some composition variation among host genera. In contrast, *Wolbachia* was sparse throughout the non-corbiculate bees. Notably, *Lactobacillus*, a major member of the Apidae-associated community, was especially negatively associated with *Wolbachia*.

### 4.1 *Wolbachia* and host-specific core taxa drive phylosymbiosis in bees

Phylosymbiosis has been studied in the corbiculate clade, which exhibits a strict and clearly resolved phylosymbiotic pattern [3]. Notably, we found that this pattern is not widely distributed across the bee tree of life, but is rather largely associated with the Apidae family. The bacterial genera defining the microbiome of the corbiculate bees – including *Bombilactobacillus, Bifidobacterium, Bombella, Gilliamella, Snodgrassella, Commensalibacter, Apibacter -* also occured in the non-corbiculate bees from the *Xylocopa* and *Centris* genera within the Apidae family, although the community composition varied among host genera. This extends previous results from Kwong et al. [3] on the core microbiome of corbiculate bees, as well as the genus-level specificity reported in Neotropical bees [5]. Because both studies focused only on corbiculate bees, they could not establish how broadly this characteristic bacterial assemblage extends across the bee phylogeny. Our results show that outside Apidae, the microbial community composition was much more variable.

Megachilidae bees, for example, largely lack most of the bacterial genera characteristic of Apidae, and instead carry a set of environmentally derived bacterial taxa, including *Sphingomonas, Corynebacterium, Methylobacterium, Stenotrophomonas,* and *Massilia*. These taxa also occur across other bee lineages, which is consistent with repeated acquisition from nest materials or floral diets maintaining a stable assemblage without social transmission [38, 39].

The distribution of the maternally inherited bacterium *Wolbachia* across the bee phylogeny showed a contrasting distribution to the Apidae microbial core. *Wolbachia* was found at very high prevalence across Halictidae, Colletidae, and Andrenidae, frequently in Megachilidae, and substantially more often in non-corbiculate Apidae. In contrast, in corbiculate Apidae, *Wolbachia* was limited to a few records in *Bombus* and reached a prevalence of only 3.8% in *Apis* species. This result is not correlated to differences in the tissues sampled in the different studies (Figure S6). Previous studies have already demonstrated that the prevalence of *Wolbachia* was higher in solitary bee tribes than in the corbiculate bees [40]. By inducing different reproductive manipulation or by affecting key life-histories in its host, Wolbachia has been described as a powerful driver of insect evolution. In bees, the bacterium has for example played a key role in the diversification of Andrena mining bees [41–44].

In contrast, after identifying a *Wolbachia*-like gene integrated within the genomes of *B. terrestris* (#LOC105666162) and 15 other *Bombus* species, Colgan et al. (2022) [43] suggested that this gene might be involved in blocking infection with the symbiont in these bee species. Unfortunately, without analyzing the host genomic material across all bee species, we could not further test whether the prevalence of this gene across bees could explain the prevalence of *Wolbachia* infection beyond the *Bombus* clade investigated by Colgan et al. (2022) [58].

Our network analyses further revealed that *Wolbachia* was negatively associated with several Apidae-associated bacteria. However, *Wolbachia* was only negatively associated with *Lactobacillus* in both corbiculate and non-corbiculate bees, after host-species block bootstrap robust test. This pattern has rarely been reported in bees, though few comparable antagonism were previously documented in *Drosophila* [45]. Host families thus structure both the presence of the core assemblage and the distribution of *Wolbachia*, and associations between them observed across lineages will need testing within hosts before they can be read as interaction.

### 4.2 Behavioral traits explain little microbiome variation beyond host phylogeny

Bees originated approximately 130 million years ago [46], and have since evolved a remarkable diversity of behavioral traits. Many of these traits show phylogenetic structure, although some, such as sociality, have evolved independently multiple times [47]. Because behavioral traits can covary with bee phylogeny, we accounted for host phylogeny when assessing their effects on the gut microbiome of bees. Conditionally on phylogeny, behavioral traits explained little of the variation in the alpha or beta diversity of the bee-associated microbial communities.

Although 90% of all bees are solitary [48], sociality is perhaps the most famous behavior of bees. Sociality explained relatively small fractions of variation in the microbiota alpha diversity. However, the advanced eusociality has arisen essentially once in bees and is confined to the corbiculate clade [49]. With so few independent contrasts available, it remains difficult to separate the effect of host phylogeny from that of the evolution of sociality. In Halictidae, a bee family characterized by social polymorphism, primitively eusocial and solitary bee species have been reported to harbor similar microbial communities [9]. Similarly, *Xylocopa* bees, which are not eusocial, were previously shown to carry a microbiota more similar to that of *Apis* and *Bombus* than to that of other solitary bees. Our analysis supports this observation, with *Xylocopa* clustering with Apidae [50] rather than with behaviorally similar taxa. In contrast, stingless bees from the families *Austroplebeia*, *Lepidotrigona* and *Tetragonisca* are all advanced eusocial taxa, and their microbial alpha diversity varies more widely than that of any other advanced eusocial taxa (Figure 2A). Together, these observations suggest that the apparent association between sociality and the bee microbiome is difficult to disentangle from host evolutionary history.

For the other behavioral traits, our findings also differ from previous studies. Although previous studies have suggested that nesting behavior affect the larval microbiome of solitary bees [51, 52], and diet has an effect on gut community composition in bees [53], we found no detectable effect from these traits on the composition of the bee microbiome. Our findings therefore suggest that the influence of these behavioral traits on the bee microbiome may be context-dependent and may not be apparent at the broader taxonomic and life-history scales we considered here. Several aspects of our study may contribute to these differences. We examined adult bees, whereas previous studies of nesting behavior focused on larvae [51, 52], and we lacked information on the specific floral or dietary resources used by individual bees. In addition, we evaluated community composition at the bacterial genus level, potentially obscuring effects that might occur at finer taxonomic resolution.

Overall, these results indicate that broad behavioral categories do not consistently predict the composition of the adult bee-associated microbiome. Instead, host phylogeny appears to provide a stronger framework for understanding microbiome variation across bees, while behavioral effects may be restricted to more specific biological scales, including during larval development, within colonies, in association with particular dietary resources, or at the microbial strain level.

### 4.2 The effect of climate differs between bee genera

Climate can affect food resources and environmental microbes, thus changing the bee microbial communities [54]. We found that the bee genera collected from arid and semi-arid regions carried more homogeneous microbiomes than those from any other climatic zone. Shifts in the composition of microbial communities due to dry climate have been observed in soils [55], and in animals [56]. For example, in the wild butterfly *Melitaea cinxia*, an extreme drought event resulted in a directional shift in the microbiome associated with caterpillars [57]. In both bees and butterflies, the patterns are consistent with narrower and more strongly filtered pools of environmental bacteria in drier habitats [58, 59], or with thermal filtering of heat-sensitive symbionts in these species [54, 60]. Despite samples spanning most climatic zones, the absence of an effect of climate on the microbiome of *Apis* bees, may further indicate that the strength of climatic effects depends on how the host acquires its microbes. Most *Apis* samples derive from managed colonies, where conserved social transmission of the core microbiota may reduce the influence of environmental variation [61, 62]. Nonetheless, our study of climatic effects on the bee microbiome is still limited by the available data, as most species occur within a single climatic zone, requiring us to pool samples within genera. Therefore, more localized climatic effects may have remained undetected.

## 5. Conclusion

Across our datasets, host phylogeny was the strongest explanatory variable for variation in the bee microbiome. Broad behavioral traits – sociality, diet or nesting preference - and climate explained little additional variation after accounting for host phylogeny, although in some bee genera, microbiomes appeared more homogeneous in arid and semi-arid regions. Phylosymbiosis was mostly observed in the Apidae-associated bacterial assemblage, and the contrasting distribution of *Wolbachia* in other bee clades, suggesting a microbial boundary at the host-family level rather than at the origin of advanced eusocial behaviors. The remarkable negative association between *Wolbachia* and *Lactobacillus* requires further investigations, as it does not yet demonstrate direct interaction between the two bacterial taxa.

Our synthesis of the bee microbiome also highlights major gaps in the field. Amongst these, a strong sampling bias towards non-corbiculate bees, and incomplete and sometimes inconsistent bee trait information, as well as missing key information in the metadata submitted to public repositories, are constraining the integration and re-utilization of public bee data. While our findings indicate that bee microbiome is shaped primarily by host lineage, we acknowledge that behavioral and climatic effects might only become visible at finer biological scales. For further synthesis work, broader taxonomic coverage and standardized reporting are essential.

## Acknowledgements

We would like to thank Mikko Kivikoski and Tobin Hammer for suggestions and comments on the study. We wish to acknowledge CSC – IT Center for Science, Finland, for computational resources. The study was funded by the Research Council of Finland (grants # 321543 and #355152) to AD, and a HiLIFE Fellowship to AD.

## Data availability statement

No new sequencing data were generated in this study. Details of the publicly available datasets used in this study, along with the analysis code, are available at https://github.com/LinyangSun/GlobalBeePhylosymbiosis.

